# Evolution of the nitric oxide synthase family in vertebrates and novel insights in gill development

**DOI:** 10.1101/2021.06.14.448362

**Authors:** Giovanni Annona, Iori Sato, Juan Pascual-Anaya, Ingo Braasch, Randal Voss, Jan Stundl, Vladimir Soukup, Shigeru Kuratani, John H. Postlethwait, Salvatore D’Aniello

**Affiliations:** Biology and Evolution of Marine Organisms, Stazione Zoologica Anton Dohrn, 80121, Napoli, Italy; Laboratory for Evolutionary Morphology, RIKEN Center for Biosystems Dynamics Research (BDR), Kobe, 650-0047, Japan; Evolutionary Morphology Laboratory, RIKEN Cluster for Pioneering Research (CPR), 2-2-3 Minatojima-minami, Chuo-ku, Kobe, Hyogo, 650-0047, Japan; Department of Integrative Biology and Program in Ecology, Evolution & Behavior (EEB), Michigan State University, East Lansing, MI 48824, USA; Department of Neuroscience, Spinal Cord and Brain Injury Research Center, and Ambystoma Genetic Stock Center, University of Kentucky, Lexington, Kentucky, USA; Department of Zoology, Faculty of Science, Charles University in Prague, Prague, Czech Republic; Division of Biology and Biological Engineering, California Institute of Technology, Pasadena, CA, USA; South Bohemian Research Center of Aquaculture and Biodiversity of Hydrocenoses, Faculty of Fisheries and Protection of Waters, University of South Bohemia in Ceske Budejovice, Vodnany, Czech Republic; Institute of Neuroscience, University of Oregon, Eugene, OR 97403, USA

**Keywords:** Vertebrate evolution, Genome duplication, Gene duplication and loss, NOS, Phylogenomics, Synteny

## Abstract

Nitric oxide (NO) is an ancestral key signaling molecule essential for life and has enormous versatility in biological systems, including cardiovascular homeostasis, neurotransmission, and immunity. Although our knowledge of nitric oxide synthases (Nos), the enzymes that synthesize NO *in vivo*, is substantial, the origin of a large and diversified repertoire of *nos* gene orthologs in fish with respect to tetrapods remains a puzzle. The recent identification of *nos3* in the ray-finned fish spotted gar, which was considered lost in the ray-finned fish lineage, changed this perspective. This prompted us to explore *nos* gene evolution and expression in depth, surveying vertebrate species representing key evolutionary nodes. This study provides noteworthy findings: first, *nos2* experienced several lineage-specific gene duplications and losses. Second, *nos3* was found to be lost independently in two different teleost lineages, Elopomorpha and Clupeocephala. Third, the expression of at least one *nos* paralog in the gills of developing shark, bichir, sturgeon, and gar but not in arctic lamprey, suggest that *nos* expression in this organ likely arose in the last common ancestor of gnathostomes. These results provide a framework for continuing research on *nos* genes’ roles, highlighting subfunctionalization and reciprocal loss of function that occurred in different lineages during vertebrate genome duplications.

## Introduction

Originally classified as a pollutant, nitric oxide (NO) was recognized as “Molecule of the Year” in 1992 [1] when its important role as a cellular signaling molecule was recognized. NO plays a role in a myriad of physiological processes, such as cardiovascular homeostasis [2], neurotransmission [3], immune response [4], and in pathological conditions such as neurodegenerative diseases [5] and cancer [6].

Nitric oxide synthase (Nos), the enzyme catalysing the biosynthesis of NO *in vivo*, is ubiquitous among organisms, including protists and bacteria [7,8]. Three *nos* gene paralogs have been described in vertebrates: two constitutively expressed genes, including *nos1* (also known as *neuronal nos*, or *nNos*), which represents the predominant source of NO involved in neurogenesis and neurotransmission [9,10], and *nos3* (*endothelial nos* or *eNos*) implicated in angiogenesis and blood pressure control in vascular endothelial cells [11,12]. In addition, *nos2* (*inducible nos* or *iNos*), which expression is instead evoked by proinflammatory cytokines, is promptly activated in a range of acute stress responses [13].

Although the availability of current genomic data covers all major ray-finned fish lineages, the evolutionary history of their *nos* gene repertoire remains puzzling. Previous studies reported a variable number of *nos* genes in teleost fishes: *nos1* is always present in a single copy; *nos2* either in one or two copies, probably due to the additional teleost-specific whole-genome duplication (TGD) [14–17], or absent as observed in several species. On the other hand, *nos3* has been reported as missing in the genomes of ray-finned fish. This apparent gene loss contrasts with literature describing a putative Nos3*-* like protein localized by antibody stains in gills and vascular endothelium of several teleost species [18,19]. The discovery of a *nos3* ortholog in the spotted gar *Lepisosteus oculatus,* a holostean fish (the sister group of teleosts within the ray-finned lineage) [20], and the variable number of teleost *nos2* genes raises new questions about the evolution of this important gene family, including: *i*. is our current view on the origin and evolution of *nos* gene family in vertebrates accurate?; and *ii*. can further investigation of *nos* expression pattern in fish retaining a *nos3* copy reveal novel functional insights? In an attempt to answer these questions, we have studied the Nos family repertoire at unprecented phylogenetic resolution, investigated conserved syntenies in fish genomes, and studied the expression pattern of all three *nos* genes during development in multiple species representing key nodes in vertebrate evolution.

## Results

### Revised evolutionary history of Nos2 and Nos3

Gaps in our current knowledge of Nos family evolution include the time of origin of the three distinct paralogous *nos* genes and when some of them were secondarily lost in specific lineages. Using sequences retrieved from public genomic and transcriptomic databases, we reconstructed a Nos phylogeny using 108 protein sequences from 53 species (see Supplementary Table 1). Species were chosen to provide a broad representation of aquatic vertebrates: cyclostomes (modern jawless fish), chondrichthyans (cartilaginous fish), and osteichthyes (bony fish) including ray- and lobe-finned fishes. Lobe-finned fishes include coelacanths, lungfishes, and tetrapods; Ray-finned fishes comprise the non-teleost lineages of polypteriformes (e.g. bichir), acipenseriformes (e.g. sterlet sturgeon), holosteans (lepisosteiformes, e.g. spotted gar, and amiiformes, e.g. bowfin), and the teleosts, subdivided into three major living lineages: osteoglossomorphs (e.g. arowana, mooneyes, and the freshwater elephantfish), elopomorphs (e.g. eels and relatives) and clupeocephalans (e.g. zebrafish and medaka) (see [21] for a recent phylogeny of ray-finned fishes).

Our phylogenetic analysis confirmed that Nos1 is present in all species of jawed vertebrates examined (Fig. 1a, green shading). In contrast, most fish lineages retained Nos2, including chondrichthyans (*Callorhinchus milii, Rhincodon typus, Chiloscyllium punctatum,* and *Scyliorhinus torazame*), polypteriformes (*Polypterus senegalus*, *Erpetoichthys calabaricus*), acipenseriformes (*Acipenser ruthenus*), holosteans (*Amia calva, Lepisosteus oculatus*), elopomorphs (*Megalops cyprinoides*), osteoglossomorphs (*Paramormyrops kingsleyae, Scleropages formosus*) and coelacanthiformes (*Latimeria chalumnae*) (Fig. 1a, grey shading), although a *nos2* gene loss event occurred at the stem of Neoteleostei (Fig. 1b), since this gene has not been found in any available genome from this clade. On the other hand, our phylogenetic analysis highlights the occurrence of extra *nos2* duplicates in several lineages, for which we adopted a specific nomenclature: in the zebrafish *Danio rerio* there are two *nos2* genes, *nos2a* and *nos2b*, while in the goldfish *Carassius auratus*, the blind golden-line barbel *Sinocyclocheilus anshuiensis* and the common carp *Cyprinus carpio* we found three: *nos2a*, *nos2ba,* and *nos2bb*; in salmonids (*Salmo salar* and *Oncorhynchus mykiss*) there are two different copies of *nos2*, named *nos2α* and *nos2β*; and last, we named *nos2.1* and *nos2.2* the two *nos2* paralogs that we found in a characid (the Mexican tetra *Astyanax mexicanus*), a gymnotid (the electric eel *Electrophorus electricus*), an ictalurid (the channel catfish *Ictalurus punctatus*), an esocid (the northern pike *Esox lucius*), and a clupeid (the Atlantic herring *Clupea harengus*) (Fig. 1a, grey shading). Our nomenclature is based both on the phylogenetic analysis and a synteny conservation analysis (see below and in the Discussion section).

**Figure 1.**
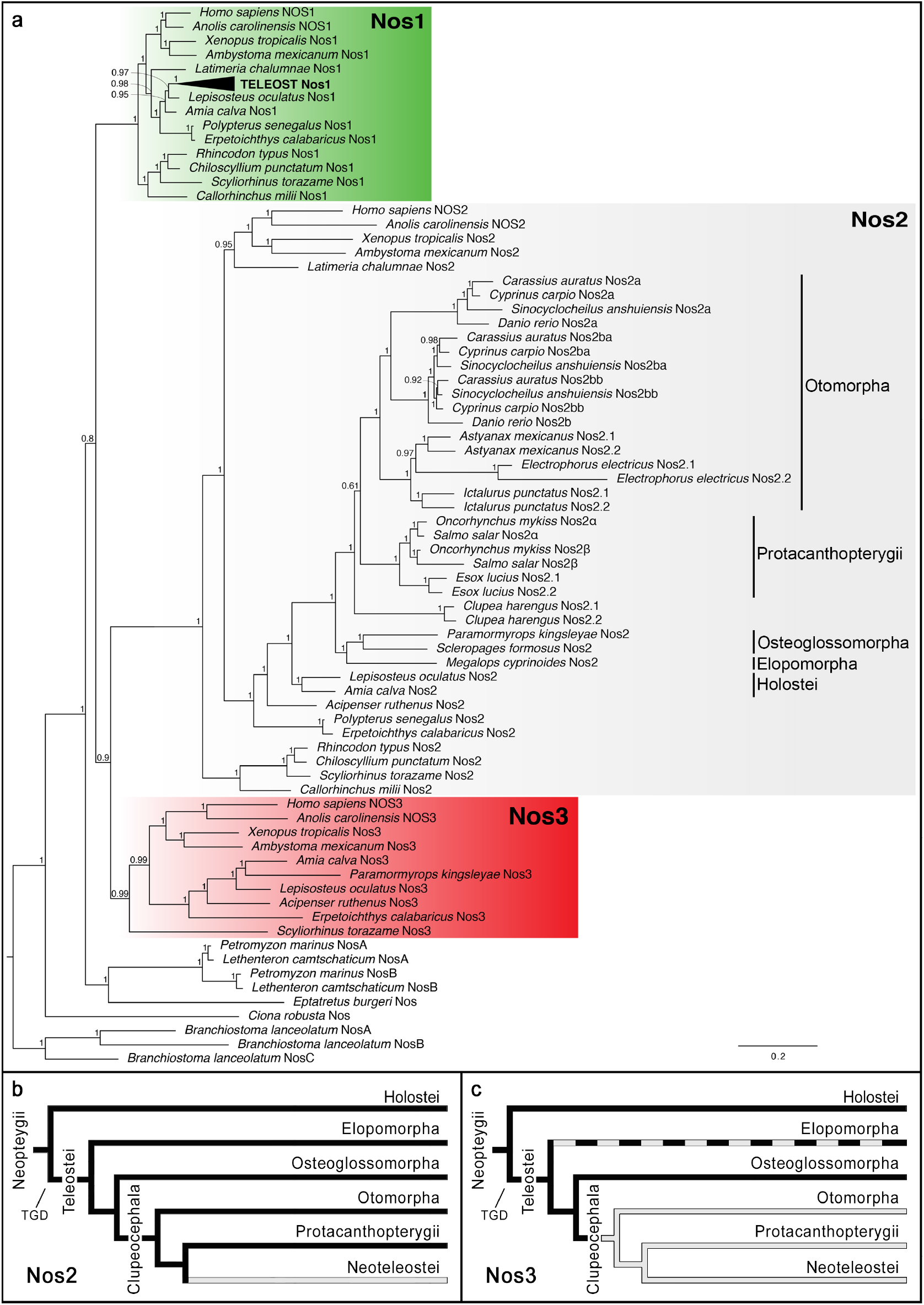
Evolution of the Nos gene family. **a**, Bayesian inference phylogenetic analysis of Nos proteins in chordates. Nos1 clade is indicated by a green shading; light grey shading for the Nos2 clade, and red shading for the Nos3 clade. Numbers at nodes represent posterior probability values. Nos proteins from invertebrate chordates, the lancelet *Branchiostoma lanceolatum,* and the tunicate *Ciona robusta*, were used as outgroup sequences. **b-c**, Evolutionary scenarios indicating the loss of Nos2 event in Neoteleostei (**b**) and Nos3 in Clupeocephala (**c**) as grey lines. Nos3 in Elopomorpha is absent although parsimony suggests it was present in stem elopomorphs, and it is indicated with a dashed line. The schematic representation of Nos1 was omitted because it is present in single copy in all analysed gnathostome species. TGD stands for Teleost-specific Genome Duplication.

Nos3 deserves special attention since it was previously believed that a loss event predated the lineage of actinopterygians or alternatively that it represents an innovation of tetrapods [8]. Nevertheless, this latter hypothesis may have been overinterpreted since few ray-finned genome sequences were originally available. The only actinopterygian *nos3* gene reported thus far is in the spotted gar [20]. Here we report the identification of *nos3* genes in genomes of the bichir *P. senegalus*, the sterlet sturgeon *A. ruthenus* [22], the bowfin *A. calva* [23], and the freshwater elephantfish *P. kingsleyae* [24] (Fig. 1a, red shading). The absence of *nos3* in all available clupeocephalans indicates a gene loss event in the stem of this group (Fig. 1c). Furthermore, we did not find *nos3* in the tarpon *M. cyprinoides*, the most complete genome available among Elopomorpha, nor in transcriptomic data of the European eel *Anguilla anguilla*. On the other hand, we did identify a *nos3* ortholog in the cloudy catshark *S. torazame,* suggesting its presence in the ancestor of gnathostomes. Previously, two *nos* genes had been found in the lamprey, called *nosA* and *nosB* [8], with unresolved orthology to gnathostome *nos1-nos2-nos3*, and derived from a lineage-specific tandem duplication in the lamprey lineage. Based on this finding, we searched for the presence of *nos* genes in other cyclostomes. In the genome of the arctic lamprey *Lethenteron camtschaticum* [25] we found orthologous genes to *P. marinus nosA* and *nosB* paralogs. On the other hand, in the inshore hagfish *Eptatretus burgeri* we identified a single *nos* gene. Our phylogenetic analysis shows that the hagfish Nos remains outside lamprey NosA-NosB clade, therefore with no clear orthology relationship to any specific gnathostome Nos1, Nos2, Nos3, and suggesting that the duplication giving rise to the lamprey *nosA-nosB* occurred at least before the last common ancestor of Petromyzontidae.

In order to better understand the gene loss and expansion events depicted by our phylogenetic analysis, we next analysed the microsynteny (genes linked in proximity) of *nos* genes in different species. This revealed a complex evolutionary scenario for *nos2* compared to *nos1* and *nos3*. Specific *nos2* duplications in different lineages are explained by distinct evolutionary events in teleosts (Fig. 2a and Supplementary Figure 1). First, the lack of synteny conservation between *nos2a* and *nos2b* in cyprinids, and the lack of *nos2a* in the expected location in non-cyprinid fishes (Supplementary Figure 1) indicates that these paralogs originated in a specific gene duplication event in a common ancestor of the lineage, independently from the TGD (the alternative explanation would require numerous *nos2a* losses in several fish lineages), in which while *nos2b* has remained the ancestral genomic location, *nos2a* has been translocated to a different position in the genome (Fig. 2a and Supplementary Fig. 1). Second, an additional genome duplication event after the TGD specifically occurred independently in several teleost lineages, causing the presence of extra *nos2* paralogs. These include some cyprinids, in which a carp-specific genome duplication event (Cs4R) likely occurred before the divergence of *C. auratus*, *S. anshuiensis* and *C. carpio* [26], and salmonids (salmonid-specific genome duplication or Ss4R) [27,28], with *S. salar* and *O. mykiss* in this study. These additional tetraploidization events can explain the origin of the two independent sets of *nos2* genes in cyprinid and salmonid species. In the case of cyprinids, both our phylogenetic and synteny analyses clearly show their *nos2b* orthology, and we denote them as *nos2ba* and *nos2bb* (Fig. 1a and Fig. 2a). In the case of salmonids, we name them *nos2α* and *nos2β* to distinguish them from the cyprinid *nos2a* and *nos2b* paralogs, which have a separate origin (see above; Fig. 2a). Third, independent tandem gene duplications explain the presence of two nos2 copies, that we named *nos2.1* and *nos2.2*, located next to each other in the same chromosomal fragment in the genomes of the Atlantic herring (*C. harengus*), the Mexican tetra (cavefish, *A. mexicanus*), the electric eel (*E. electricus*), the channel catfish (*I. punctatus*), and the northern pike (*E. lucius*) (Fig. 2a).

**Figure 2.**
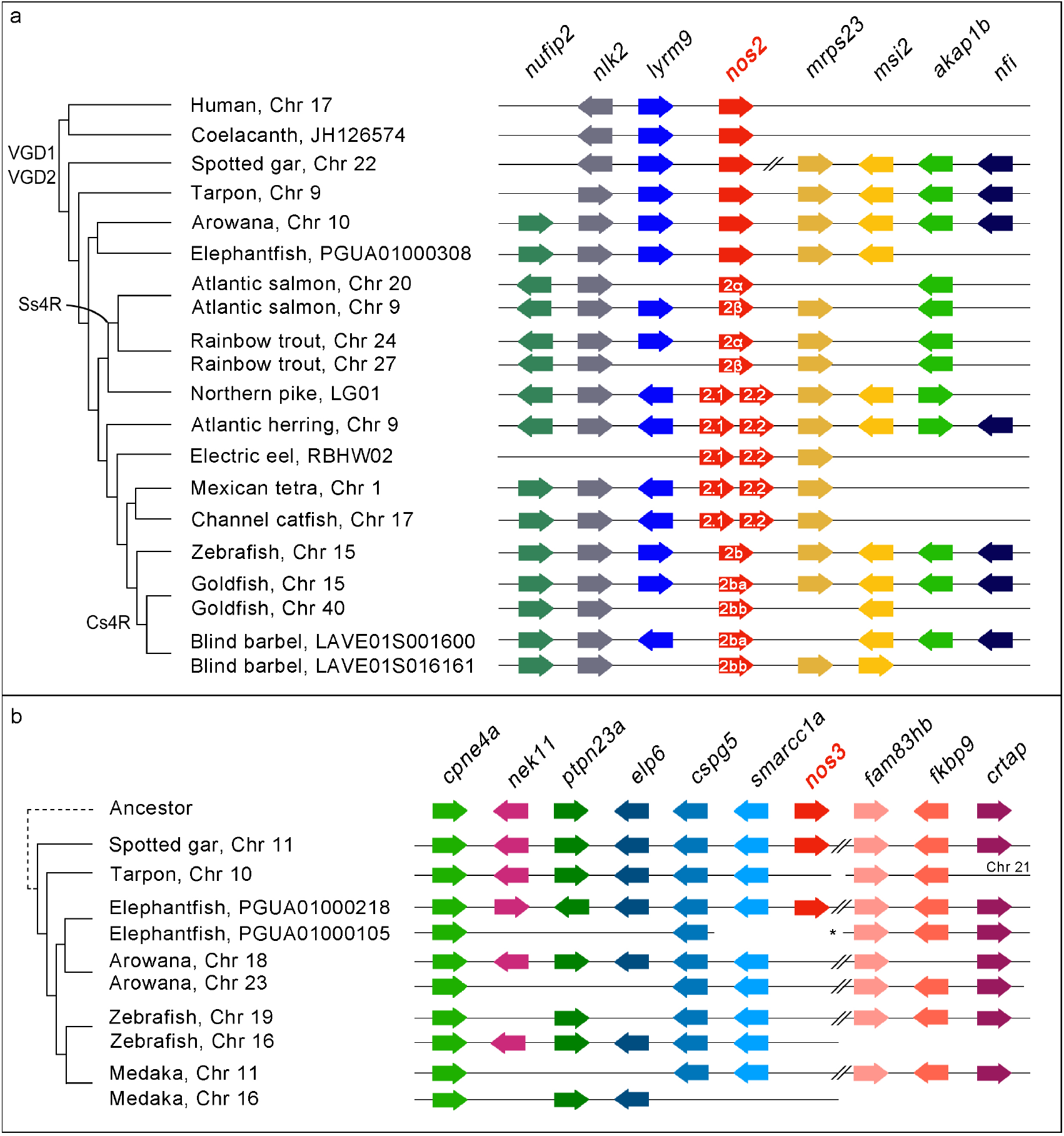
Conserved microsynteny of *nos2* and *nos3.* **a**, The *nos2* paralogs derived from different duplication modalities: carp-specific genome duplication (Cs4R) (*nos2ba* and *nos2bb* in the goldfish and blind golden-line barbel); salmonid-specific genome duplication (Ss4R) (*nos2α* and *nos2β* in the Atlantic salmon and rainbow trout); tandem gene duplication occurred independently in five lineages (*nos2.1* and *nos2.2* in the northern pike, Atlantic herring, electric eel, Mexican tetra and channel catfish). An additional *nos2* duplicate (*nos2a*) is present in cyprinids (zebrafish, goldfish, and blind barbel). **b**, A conserved synteny map of genomic regions around the *nos3* gene locus highlights the loss in Clupeocephala (including zebrafish and medaka), and in Osteoglossomorpha (arowana). Genes are represented as arrows and are coulor coded according to their orthology and ohnology. The direction of arrows indicates transcription orientation. The symbol // indicates a long-distance on a chromosome. The asterisk indicates scaffold 72 of the freshwater elephantfish genome [24].

Bichir, reedfish, sterlet, spotted gar, bowfin and freshwater elephantfish are the only ray-finned fishes that retained a *nos3* ortholog. Therefore, we investigated the absence of *nos3* in clupeocephalans. First, we looked for the genomic region containing *nos3* in fishes that represent outgroups to the clupeocephalans. We found one long scaffold of the *P. kingsleyae* genome (scaffold 217) [24] showing extensive conserved synteny with the *nos3*-containing segment of the linkage group 11 (LG) in the spotted gar genome (Fig. 2b). While these appear to correspond to one of the TGD ohnologons (Fig. 2b), there are other two *P. kingsleyae* scaffold segments (from scaffolds 72 and 104) that together seem to represent the second TGD ohnologon, but lacking the expected *nos3* TGD ohnolog (Fig. 2b). Zebrafish chromosomes 16 and 19 and medaka chromosomes 11 and 16 contain orthologous regions to the two *P. kingsleyae* and *L. oculatus* TGD ohnologons, but lack a *nos3* gene at the expected locations. One-to-one relationship between these *P. kingsleyae* scaffolds and zebrafish and medaka chromosomes is challenging to determine (Fig. 2b). Regardless, the most parsimonious explanation for the nos3 repertoire in ray-finned fishes is that, first, one of the two *nos3* TGD ohnologs was lost in the teleost common ancestor, while the other was retained and later lost in secondary, independent events in the common ancestor of Clupeocephala and, probably, that of Elopomorpha (Fig. 1c and Fig. 2b).

### Expression of *nos* in vertebrate developing gills

Spotted gar is an important emerging model organism because it represents an evolutionary bridge between teleosts and tetrapods that facilitates cross-species comparisons. The gar genome is slowly evolving compared to that of teleosts and has preserved a more ancient structural organization [29]. Therefore, we examined the expression patterns of *nos* genes during gar development. As expected, based from the literature, *nos1* was expressed in several regions of the developing nervous system (Supplementary Fig. 2). In contrast, *nos2* expression was not detected during the developmental stages covered in the present study, i.e., from 4 to 14 days post fertilization (dpf). Unexpectedly, the expression of *nos3* was first detected in gar embryos in the pharyngeal area at 4 dpf (Fig. 3a-b) and increased at 6 dpf (Fig. 3c-d). At 7 dpf, embryos showed clear *nos3* expression in developing arches III, IV, and V (Fig. 3e-g). Later, at 11 dpf, the positive signal is localized in gill filaments (Fig. 3i-k). Histological sections highlighted the presence of *nos3* in the epithelium of branchial lamellae (Fig. 3l), also confirmed by the signal in gill structures in an advanced stage of maturation in 14 dpf juveniles (Fig. 3m-p).

**Figure 3.**
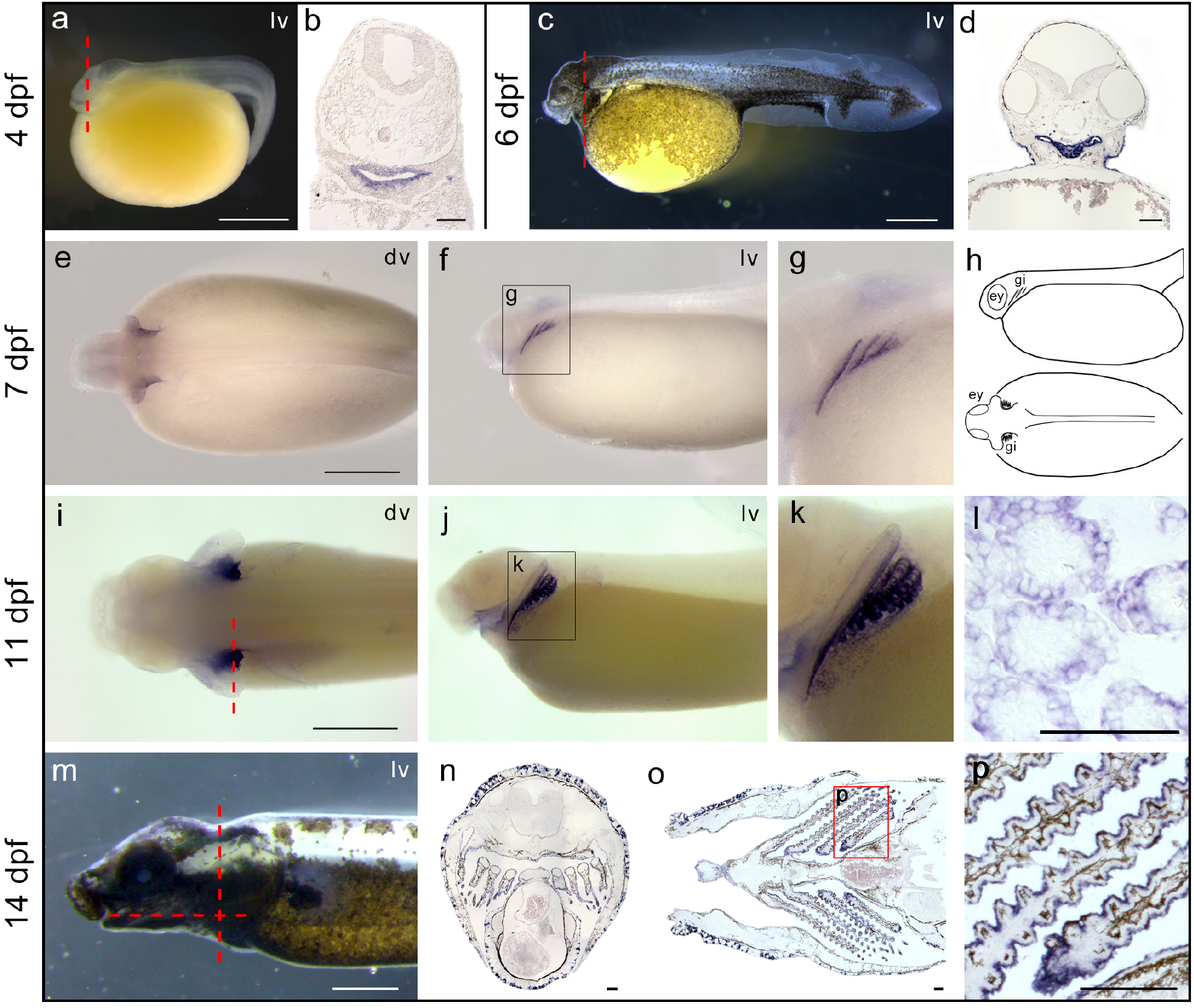
Spotted gar *nos3* localization during development. Expression of *nos3* is localized in the pharyngeal area in 4 dpf (**a-b**) and 6 dpf (**c-d**) embryos, in pharyngeal arches in 7 dpf larvae (**e-g**) schematized in (**h**), in developing gills in 11 dpf late larvae (**i-l**), and in gill lamellae in 14 dpf juveniles (**m-p**). Coronal (**n**) and transversal section (**o**) planes are indicated with a red dashed line in (**m**). Abbreviations: ey, eye; gi, gill; dv, dorsal view; lv, lateral view. Scale bar is 1 mm in a, c, e, i, m; 100 μm in b, d, l, n, o, p.

The detection of *nos3* transcripts in gills of spotted gar and the established involvement of NO gas in osmoregulatory control and vascular motility in gills of numerous teleosts [30–35] prompted us to investigate whether a similar *nos* expression patterns occurred in developing gills of other fish species. We investigated *nos* expression in the sterlet sturgeon *A. ruthenus* and the bichir *P. senegalus,* members of early-branching groups of ray-finned fishes [21]. Moreover, we similarly investigated *nos* expression in the chondrichthyan cloudy catshark *S. torazame* to infer the ancestral expression condition among gnathostomes. Unlike gar, we discovered that *nos3* was not expressed in gills of other species analysed in this work (Supplementary Fig. 2), thus raising questions about whether *nos3* expression in gills represents an oddity of holosteans or gars. Surprisingly, *nos1* and *nos2* were expressed in gills of sturgeon, bichir, and shark. In particular, *nos2* was expressed in the branchial area of the sterlet sturgeon (Fig. 4a-c) and bichir embryos (Fig. 4d-f), while *nos1* is expressed in gills of catshark embryos (Fig. 4g-i).

**Figure 4.**
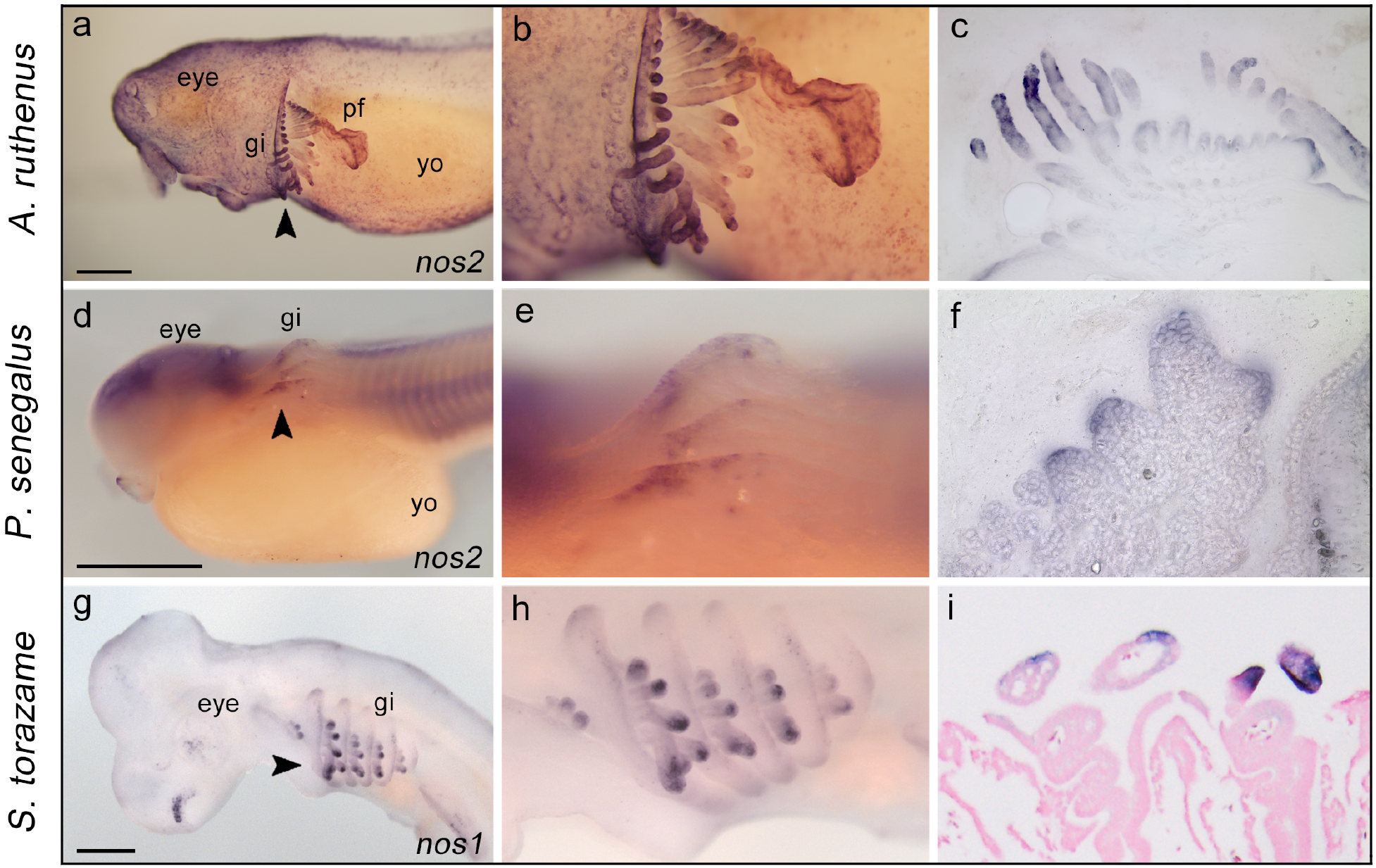
Expression of *nos* genes in developing gills of sterlet sturgeon, bichir, and shark embryos. The expression of *nos2* in the gills of sterlet sturgeon *Acipenser ruthenus* (14 mm stage, **a-c**) and bichir *Polypterus senegalus* (stage 31, **d-e**); *nos1* in the shark *Scyliorhinus torazame* (stage 27, **g-i**). Higher magnification views of the gill structure of **a, d, g** are shown in **b, e**, **h**, respectively. The arrowheads indicate sectioning plane (**a, d, g**): transversal sections (**c, f**, 50 μm) and frontal section (**I**, 10 μm). Abbreviations: gi, gill; yo, yolk; pf, pectoral fin. Scale bar in a, d, g is 0.5 mm.

Our results show that *nos* paralogs are expressed in pharyngeal arches and gills in both actinopterygians and chondrichthyans. These findings lead us to question whether *nos* expression in gills could be a conserved feature also in sarcopterygians, and in particular in amphibians that use gills for gas exchange. Therefore, to investigate the presence of *nos* transcripts in amphibia, we chose the neotenic axolotl *Ambystoma mexicanum* because it retains functional external gills throughout life. Gene expression analysis by qPCR revealed that *nos1* and *nos2* are almost not detectable in adult axolotl gills, while *nos3* turned out to be highly expressed in gill structures (Supplementary Fig. 3). Therefore, we conclude that *nos* expression in gills is a conserved feature in neotenic amphibian assayed, previously observed exclusively in fishes.

### Expression of *nos* genes in the lamprey

In cyclostomes (jawless vertebrates, including lampreys and hagfish), cartilaginous and bony gnathostomes (jawed vertebrates), gills are endoderm-derived structures, pointing to a single origin of pharyngeal gills before the divergence of these vertebrate lineages [36,37]. The two lamprey *nos* paralogs, *nosA,* and *nosB*, display an unresolved orthology relationship with their gnathostomes *nos1*, *nos2,* and *nos3* (Fig. 1). To assess whether *nosA* and *nosB* are expressed in gills during embryogenesis, we performed whole-mount *in situ* hybridization experiments at different embryonic stages. We found that lamprey *nosA* was expressed in several tissues, including the brain, dorsal midline epidermis, tailbud, mouth, and cloaca, but not in gills (Fig. 5a-b). Conversely, the lamprey *nosB* paralog showed restricted expression in the developing mouth, specifically in the cheek process, including upper and lower lip regions (Fig. 5c-d). These results show that in the arctic lamprey, neither of the two *nos* paralogs is expressed in immature or mature gills, suggesting a fundamental difference in the role of *nos* genes in jawless and jawed vertebrates.

**Figure 5.**
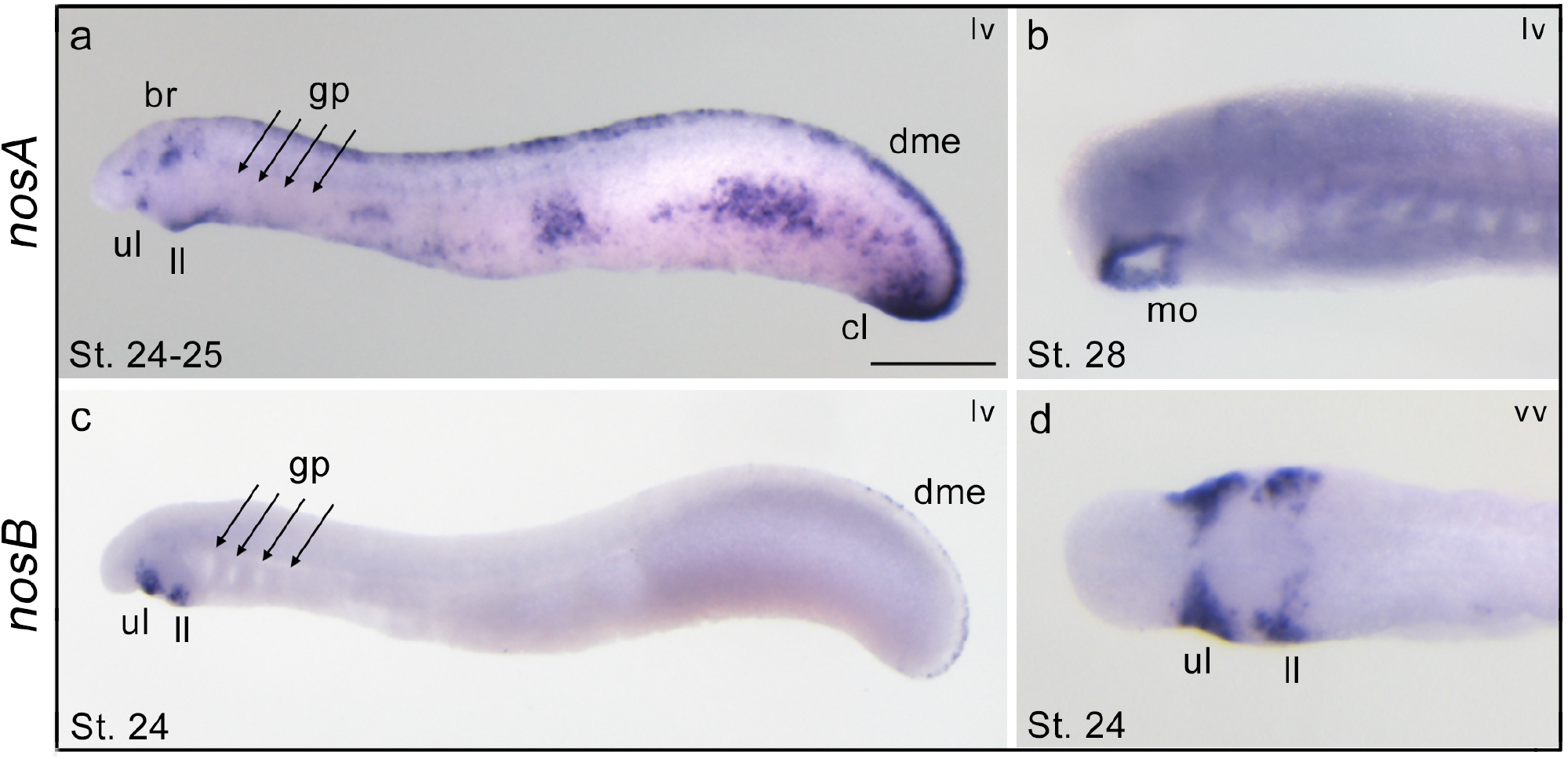
Expression patterns of *nosA* and *nosB* in larvae of the arctic lamprey. At stage 24-25 the *nosA* is expressed in the brain, mouth, upper and lower lip, dorsal midline epidermis, and cloaca (**a**). At stage 28, *nosA* expression is restricted to the mouth (**b**). The *nosB* is exclusively expressed in the cheek process, consisting of upper and lower lips (**c-d**), and faint expression in the dorsal midline epidermis (**c**). Abbreviations: br, brain; cl, cloaca; dme, dorsal midline epidermis; gp, gill pouches; mo, mouth; ll, lower lip; ul, upper lip; lv, lateral view; vv, ventral view. Scale bar in a is 0.5 mm.

## Discussion

Actinopterygian fishes experienced one of the largest radiations in the animal kingdom and their history represents a valuable resource for the formulation of hypotheses regarding the evolution of vertebrate gene families. In this work, we employed data from recent genome projects to clarify and update the evolution of Nos family across vertebrates. We performed a phylogenetic reconstruction using Nos protein sequences from key vertebrate groups, including cyclostomes for which little information has previously been available. Our phylogenetic analysis confirmed that Nos1 is ubiquitously present as single copy gene across the gnathostome lineage, at least in the covered osteichthyan and chondrichthyan species. Branch lengths of the Nos1 clade suggest a slow evolutionary rate throughout vertebrate evolution in respect to the other two *nos* genes. Furthermore, our phylogenetic data, complemented with syntenic analyses, highlighted for the first time a highly complex scenario of Nos2 evolution, for which we suggest a nomenclature that attempts to incorporate evolutionary origins into gene names. Previous analyses showed the presence of two *nos2* genes (*nos2a* and *nos2b*) in zebrafish and goldfish [38,39]. Here we show the presence of a *nos2a* paralog also in other two cyprinids, *C. carpio* and *S. anshuiensis* (Fig. 1a and Fig. 2a). *nos2a* and *nos2b* likely derive from an event of gene duplication that occurred specifically at the stem of the group, and not related to the classic TGD. This result is supported by synteny analysis since the chromosomal position of *nos2a* and *nos2b* genes is not conserved (Fig. 2a and Supplementary Fig. 1), as it would be expected if they were retained after a whole-genome duplication. On the other hand, the cyprinid *nos2b* paralog independently duplicated in carps after the Cs4R [26], as the conserved synteny suggests (Fig. 2a). In salmonids, synteny analysis also implies that the two Nos2 paralogs originated secondarily after the Ss4R (Fig. 2a) [27,28]. Here, we call these genes *nos2ba* and *nos2bb* in carps to emphasize and clarify their relationships to zebrafish genes, and *nos2α* and *nos2β* in salmonids to indicate their distinct evolutionary origin. Additionally, the present work shows that *nos2* has undergone several independent lineage-specific tandem gene duplication events (*nos2.1* and *nos2.2*) (Fig. 2a). The search of *nos2* in available fish genomes, covering all main groups, failed to find it in any Neoteleostei, and for this reason, we hypothesized a *nos2* gene loss event occurred in stem Neoteleostei (Fig. 1 and Fig. 6). It is worth mentioning that NO produced upon stimulation of the inducible *nos* (*nos2*) is considered one of the most versatile players of the immune system against infectious diseases, autoimmune processes and chronic degenerative diseases [4,40]. For this reason, it would be important in the future to investigate the impact of Nos2 loss on the immune response in Neoteleostei and if any compensatory mechanisms occurred through the activation of other *nos* paralogs. In addition, Nos2 is the only *nos* gene with retained duplicates in vertebrates, therefore, it would also be important to understand if *nos2* duplicates underwent neofunctionalization or subfunctionalization, thus providing new functional features to the organism.

**Figure 6.**
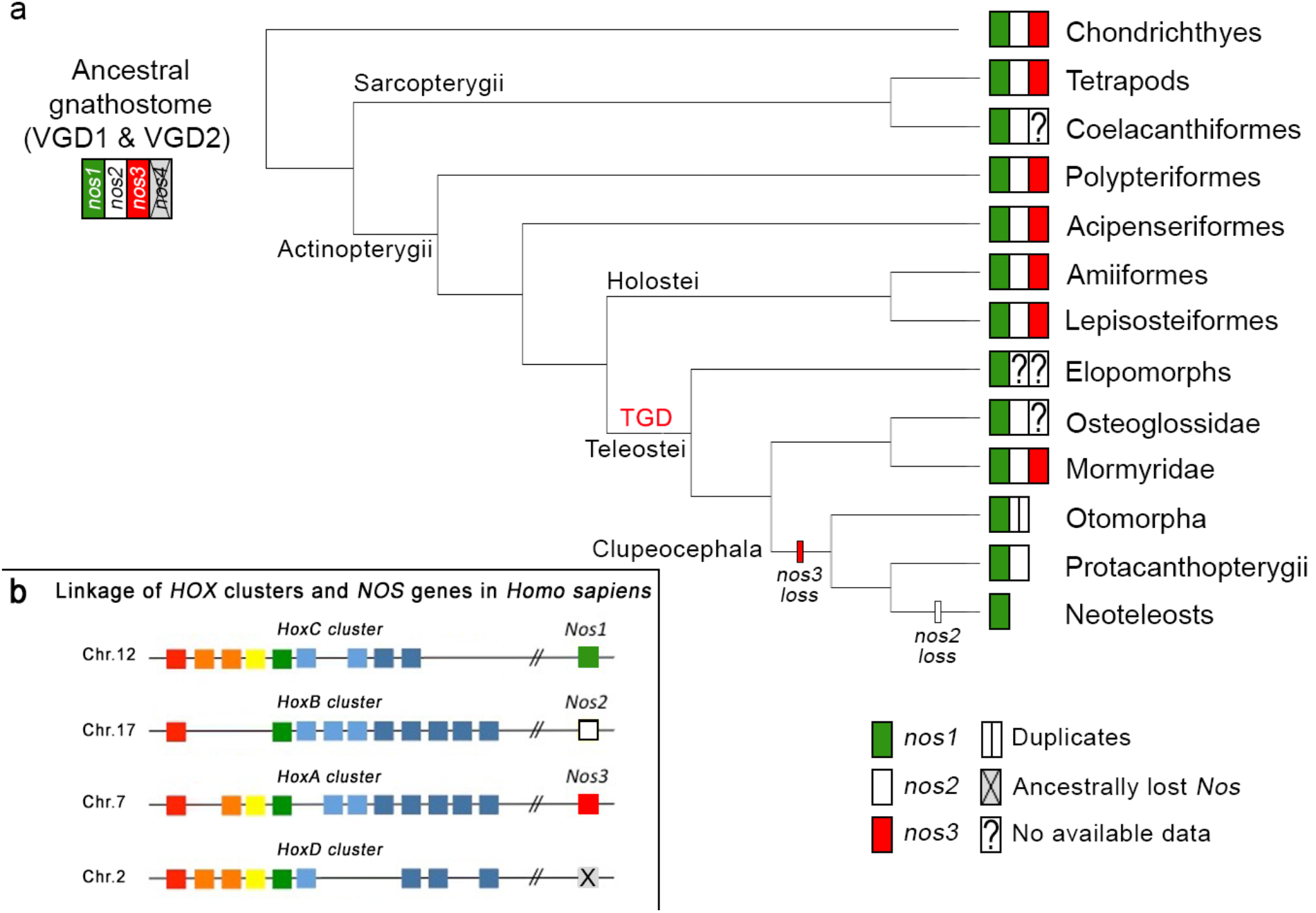
*Nos* evolution in light of recent gene findings in vertebrates. The proposed evolution of *nos* genes in gnathostomes (**a**) supposes an ancestral loss of a predicted fourth *nos* gene (grey box), based on the linkage of human *nos* and *Hox* clusters (**b**). Loss of *nos3* occurred in stem Clupeocephala and loss of *nos2* in stem Neoteleostei (**a**). Species-specific *nos2* duplications occurred in some Otomorpha, including Cyprinidae and Characidae families.

Concerning *nos3*, our understanding of its evolutionary history had a twist with the finding of a *nos3* ortholog in the spotted gar genome [20], proving that the previously postulated actinopterygian-specific loss of *nos3* was an incorrect inference. Fostered by this discovery, we specifically searched for the presence of *nos3* orthologs in a wide range of fish species to infer the ancestral condition. We identified a *nos3* gene in bowfin, thus confirming the presence of *nos3* in the other reference genus of the holostean clade, in addition to gar (Fig. 1a and Fig. 6). Furthermore, the presence of *nos3* in genomes of bichir and sterlet sturgeon, which diverged prior to the teleostean and holostean split, confirmed the hypothesis that *nos3* was already present in the common ancestor of extant osteichthyans, rather than an innovation of tetrapods [8] or neopterygians (holosteans plus teleosts) [20] (Fig. 6). We did not find *nos3* gene in the tarpon *M. cyprinoides* genome (Fig. 2b), and to date, the limited genomic and transcriptomic data of eels, congers, and morays cannot endorse the presence of a *nos3* in Elopomorpha. Therefore, more genome sequences are necessary to confirm its absence in this key group. We also did not find *nos3* in any fish from Clupeocepahala (non-elopomorph and non-osteoglossomorph teleosts; Fig. 1a and Fig. 2b) suggesting that a loss event took place in the common ancestor of clupeocephalans (Fig. 1c and Fig. 2b). Notably, we found a *nos3* gene in the osteoglossomorph elephantfish *P. kingsleyae* (Fig. 1a and Fig. 2b), and it allowed us to confirm that the loss of *nos3* did not occur in the last common teleost ancestor, as previously thought [20]. These findings suggest instead the following evolutionary scenario for the *nos3* gene: first, since we only find a maximum of one *nos3* gene in those cases where it is present, we assume that one of the two TGD ohnologs was immediately lost after the TGD, and the other one was retained. This *nos3* gene was then lost in the ancestors of elopomorphs –although further research is needed to confirm this– and clupeocephalans independently in separate events (Fig. 6).

The discovery of *nos3* in sharks (*S. torazame* in this study) suggests that the origin of *nos3* predates the divergence of gnathostomes and that three distinct *nos* paralogs were already present in the last common ancestor of gnathostomes (Fig. 6), likely originating after the two rounds of whole-genome duplication that took place during early vertebrate evolution (VGD1 and VGD2, 2R hypothesis) [41]. The origin of *nos* genes is, in fact, supported by the linkage to the evolutionarily conserved *Hox* gene clusters and several other syntenic genes (Fig. 6b and Supplementary Fig. 4). Under this scenario, then a fourth *nos* gene (putative *nos4*) should have existed but was apparently lost early in the gnathostome evolution (Fig. 6a).

The apparent lack of *nos* genes in some vertebrate lineages remains to be clarified, such as the absence of *nos3* in coelacanth *L. chalumnae* (an extant basally diverging sarcopterygian), in arowana *S. formosus* (an osteoglossomorph), and in elopomorph fishes. In the future, further genomic projects will surely fill these gaps in our understanding of this fascinating gene family.

The importance of NO in the ontogeny and function of vertebrate gills has already been documented in the context of physio-pharmacological studies, primarily using inhibitors of Nos activity. In gills, NO acts as a paracrine and endocrine vasoactive modulator and, therefore, plays a crucial role in the distribution of oxygenated blood [42]. Moreover, NO has an osmoregulatory function controlling the movement of ions across the gill epithelium [33,43–45], and represents an important molecular component of the immune system employed by macrophages to attack and destroy pathogens [46]. Nevertheless, documentation of Nos enzymatic activity in fish gills has relied exclusively upon techniques unable to discriminate among individual Nos proteins, such as NADPH-diaphorase activity and immunolocalization with heterologous mammalian antibodies [42,44,45,47]. Therefore, the detected enzymatic activity has for a long time been indicated generically as ‘Nos-like’. Here we used a different approach based on mRNA transcript detection methodology, which unequivocally distinguishes different genes, and showed, for the first time, that indeed *nos* genes are expressed in gills during development in various vertebrates. However, surprisingly different Nos paralogs are expressed in gills in different animals tested: *nos1* in shark, *nos2* in bichir and sterlet sturgeon, and *nos3* in spotted gar. The most parsimonious hypothesis to explain this result is that the ancestral *nos* gene had a number of roles in gills, immune system, brain, and other organs that was controlled by separate regulatory elements and, due to subfunctionalization after the vertebrate 2R (according to the Duplication-Degeneration-Complementation (DDC) model) [48], these physiological roles partitioned to different *nos* ohnologs as lineages diverged and reciprocal loss of the gill expression function occurred in a lineage-specific way. Further support to this hypothesis comes from the identification of *nos1*-positive cells in gill of zebrafish at 5 dpf, in addition to brain, eye, periderm and NaK ionocytes, according to the recently released developmental single-cell transcriptome atlas [49] (Supplementary Fig. 5).

Additionally, to corroborate the involvement of NO in normal gill physiology, we searched for *nos* expression in gills of a paedomorphic amphibian, the Mexican axolotl, which maintains gill structures in adulthood. Taking into account the different evolutionary and developmental origin of internal and external gills [50], the conservation of *nos3* expression in gills indicated that the NO signaling system could be indeed fundamental for the physiology and development of this structure in the axolotl, and perhaps generally in pre-metamorphic amphibians. Therefore, our data obtained from established and emerging model species highlighted that the expression of at least one *nos* gene has a functional role in gnathostome gills.

Recently, a single origin of pharyngeal gills predating the divergence of cyclostomes and gnathostomes was suggested [36]. Therefore, we investigated whether either of the two arctic lamprey *nos* paralogs is expressed in developing gills, but found them expressed mainly in the nervous system, mouth and pharynx, similarly to the expression pattern previously reported in the cephalochordate amphioxus [51,52]. Nevertheless, we found neither of the two genes to be expressed in gills during lamprey development, leading us to speculate that either the expression of *nos* genes in gills was acquired in gnathostomes after the divergence from cyclostomes, or alternatively, gill expression was a feature of their last common ancestor but lost in the lamprey lineage. Future work on hagfish embryology would be necessary to help solve this issue.

In conclusion, our findings pave the way for future studies that aim to investigate the ontogenetic role of nitric oxide in gill development of aquatic vertebrates, possibly by loss-of-function approaches using either *Nos* protein-specific chemical inhibitors or CRISPR-Cas9 gene editing. From the perspective of the evolution of developmental mechanisms, it would be interesting to understand more about the species-specific regulatory mechanisms that drive different *nos* genes expression patterns in gills in different species

## Materials and Methods

### Sequence mining and phylogenetic analysis

Nos sequences used for evolutionary analyses were retrieved from NCBI (https://www.ncbi.nlm.nih.gov/), Ensembl (www.ensembl.org/index.html), Skatebase (http://skatebase.org/) and DDBJ (https://www.ddbj.nig.ac.jp/index-e.html) databases, using direct browser webpages or by downloading fully assembled genomes and transcriptomes (see Supplementary Table 1 for accession numbers). The quality of protein sequences was checked and, where needed, manually curated excluding from the dataset partial or low blast score sequences. We used proteins from *Homo sapiens*, *Anolis carolinensis* and *Xenopus tropicalis* as internal references, and two non-vertebrate chordates as outgroups: the cephalochordate *Branchiostoma lanceolatum* NosA, NosB and NosC, and the tunicate *Ciona robusta* Nos.

Nos sequences from bowfin (*A. calva*) were obtained from a draft genome assembly [23]. Lamprey *nosA* and *nosB* genes were obtained by TBLASTN v2.2.31+ searches [53] from the v1.0 draft genome of arctic lamprey *L. camtschaticum* [25] and the germ line draft genome of sea lamprey *P. marinus* [54]. Initial predictions were extended, corrected, and confirmed by RACE PCRs in the case of *L. camtschaticum*. Both *P. marinus nosA* and *nosB* were manually curated using Wise2 [55]. The single *nos* gene sequence from the inshore hagfish *E. burgeri* was obtained from a *de novo* transcriptome assembly [56]. Sequences of *nos1*, *nos2,* and *nos3* genes from the cloudy catshark *S. torazame* were obtained from a *de novo* transcriptome assembly [56] employing TBLASTN v2.2.31+. A partial *nos1* sequence (g15096.t1) was found in the European eel *A. anguilla* transcriptome database (EeelBase 2.0, http://compgen.bio.unipd.it/eeelbase/), but it was deliberately excluded from the phylogenetic analyses because of alignment ambiguities, probably due to gene assembly errors.

For phylogenetic analysis, Nos amino acid sequences were aligned using the MUSCLE algorithm [57] as implemented in MEGAX (version 10.2.4) [58], with default parameters, run in a MacOS 11.2.1 operating system and saved in FASTA format. The alignment was trimmed by trimAl v1.2rev59 [59], using the ‘-automated1’ parameter. The trimmed alignment was then formatted into a nexus file using readAl (bundled with the trimAl package) (supplementary file S1). A Bayesian inference tree was constructed using MrBayes v3.2.6 [60], under the assumption of an LG+I+G evolutionary model. Two independent MrBayes runs of 2,000,000 generations were performed, with four chains each and a temperature parameter value of 0.05. The tree was considered to have reached convergence when the standard deviation stabilized under a value of <0.01. A burn-in of 25% of the trees was performed to generate the consensus tree (1,500,000 post-burnt-in trees).

### Synteny

Conserved synteny analyses were manually performed on fish chromosomes or scaffolds. With the aim of finding synteny blocks flanking the *nos2* and *nos3* orthologs, we employed the Synteny Database (http://syntenydb.uoregon.edu/synteny_db/) [61,62]. Additional information was retrieved in NCBI (https://www.ncbi.nlm.nih.gov/), Ensemble v.102 (http://www.ensembl.org/index.html) and Genomicus v100.01 (https://www.genomicus.biologie.ens.fr/genomicus-100.01/cgi-bin/search.pl) [61].

### Collection of embryos and tissues

Spotted gar *L. oculatus* adult specimens were collected from the Atchafalaya River basin, Louisiana (USA) and cultured in a 2 m diameter tank containing artificial spawning substrate. Spawning was induced by injection of Ovaprim© (0.5 ml/kg) and embryos were raised in fish water (salinity 1 ppt) at 24°C in a 14/10 h light/dark cycle [63]. The developmental staging was determined following hours or days post fertilization in addition to morphological criteria [64].

Embryos of *L. camtschaticum* were obtained by artificial fertilization, cultured at a temperature ranging between 9 and 12°C, and staged as previously described [65,66]. Embryos of *S. torazame* were obtained, cultured, and staged as previously described [67,68].

Bichir *P. senegalus* embryos were obtained from the breeding colony at the Department of Zoology, Charles University, Prague (Czech Republic) by natural breeding. Embryos were kept at 28°C and staged using Diedhiou and Bartsch (2009) guidelines [69]. Sterlet sturgeon *A. ruthenus* embryos were obtained from the hatcheries of the Research Institute of Fish Culture and Hydrobiology in Vodnany, University of South Bohemia (Czech Republic). Embryos were raised in tanks containing E2 Pen/Strep zebrafish medium and incubated at 17°C until the desired stages, according to Dettlaff and collaborators (1993) [70]. Gill tissues from two adult axolotls (RRID:AGSC_110A) were collected under benzocaine anesthesia (University of Kentucky, USA, IACUC protocol 2017-2580).

### Gene expression analysis by *in situ* hybridization

For all species used in the present study, total RNA was isolated from a mix of embryo stages using the phenol-chloroform method with TRIzol (Thermo-Fisher Scientific). cDNA was synthesized from 1 μg of total RNA using the SuperScript VILO cDNA Synthesis kit (Thermo-Fisher Scientific). Primers for PCR amplification are listed in Supplementary Table 2. Amplicons were cloned into the pGEM-T Easy Vector (Promega) and Sanger sequenced. Antisense Digoxygenin-UTP riboprobes were synthesized using SP6 or T7 RNA polymerases and the DIG RNA Labeling kit (Roche).

Whole-mount *in situ* hybridization experiments were performed following protocols previously described: spotted gar [71], bichir and sturgeon [72], lamprey [65], and shark [73], with slight modifications. For spotted gar embryos at 7 dpf (Long & Ballard stage 24) and 11 dpf (Long & Ballard stage 28), longer proteinase K (10 μg/mL) digestion times were performed, respectively 25 and 35 minutes at 24°C. Moreover, endogenous melanin pigment was removed using bleaching solution [(3% hydrogen peroxide (H_2_O_2_) and 1% potassium hydroxide (KOH) in distillate water (ddH_2_O)] for a few minutes. For 14 dpf gar embryos (Long & Ballard stage 31), we performed *in situ* hybridizations on cryosections, as previously described [74], including modifications reported in [75].

Transversal vibratome sections of bichir and sturgeon embryos (thickness 50 μm) were made on whole-mount hybridized embryos upon embedding in gelatin/albumin/glutaraldehyde [50]. Shark embryos were embedded in paraffin after whole-mount *in situ* hybridization assays, and frontal sections (10 μm) were obtained with a microtome.

Whole-mount and sectioned preparations mounted on slides were imaged on Axio Imager Z2 with Apotome 2 (Carl Zeiss), equipped with Axiocam 503 coulor digital camera and Axio Vision software for analysis. Whole-mount bichir and sturgeon embryos were photographed as Z-stacks using a motorized dissection microscope (Olympus SZX12) and deep-focus images were generated by merging Z-stacks in QuickPhoto Micro.

### Real-time PCR

Expression levels of *nos* genes in axolotl *A. mexicanum* gills were analysed by qPCR using specific primers reported in Supplementary Table 2. The *atpf51* gene was used as a reference. RNA was isolated by performing a chloroform extraction and isopropanol precipitation. RNA was quantified using a Nanodrop and 1 μg of total RNA was used to generate cDNA using an Invitrogen Super-Script IV cDNA synthesis kit with oligo-dT tailing. RT-qPCR was performed in triplicate with SYBER Master Mix on a LightCycler 96 (Bio-Rad), using a 2-step amputation protocol (95°C for 10 sec, 60°C for 30 sec) and 40 cycles. Data were analysed using the ΔΔCT method.

## Data availability

Accession numbers of protein sequences used in the phylogenetic analysis are available in Supplementary Table 1. Primer sequences used for the synthesis of *in situ* hybridization riboprobes and in quantitative real-time PCR experiments are given in Supplementary Table 2.

## Acknowledgments

The authors thank Allyse Ferrara and Quenton Fontenot, Louisiana State University (USA), for their help in the generation of spotted gar embryos. The authors thank Fumiaki Sugahara for his help in the interpretation of results in the arctic lamprey, Anna Pospisilova for technical assistance with bichir and sturgeon *in situ* hybridizations, and Martin Psenicka, Roman Franek, Michaela Fucikova, Marek Rodina, David Gela, and Martin Kahanec for sterlet sturgeon spawns. A special thanks to Robert Cerny for the establishment of the African bichirs colony at the Charles University in Prague.

Giovanni Annona was supported by the Research grant POR Campania FSE 2014/2020 (IT) and by the EMBO Short Term Fellowship (# 6936) to visit the Postlethwait laboratory in Oregon (USA) and for the field trip in Louisiana (USA). Jan Stundl is supported by the European Union’s Horizon 2020 research and innovation program under the Marie Skłodowska-Curie grant agreement No. 897949. Vladimir Soukup is supported by the Charles University Research Centre program No. 204069 and grant SVV260571/2020. Randal Voss and the Ambystoma Genetic Stock Center are supported by the National Institutes of Health, USA (P40OD019794). John H. Postlethwait is supported by the R01 OD011116 grant from the US National Institutes of Health. Salvatore D’Aniello is supported by the NOEVO grant from the Stazione Zoologica Anton Dohrn Napoli.

