## Additional Information for "Evolution of the nitric oxide synthase family in vertebrates and novel insights in gill development"

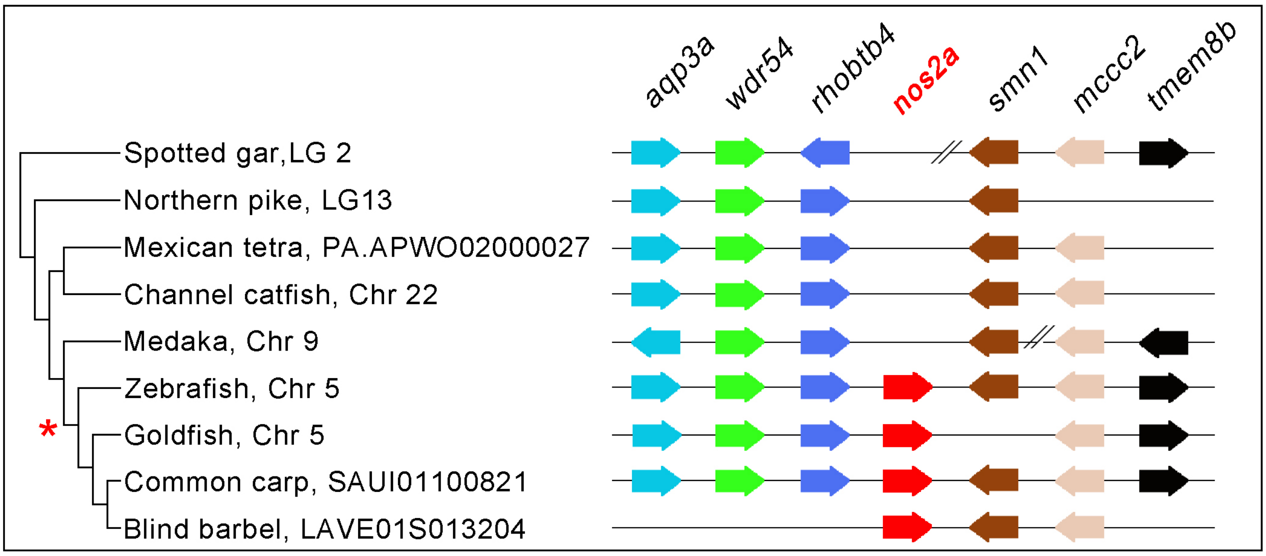


**Supplementary Figure 1.** Synteny conservation analysis of *nos2a* genes. Orthologs from different species are represented with the same colour code. The direction of arrows indicates chromosomal gene orientation. The symbol // indicates long distance on the chromosome. The red asterisk indicates that *nos2a* has been gained exclusively in cyprinids.

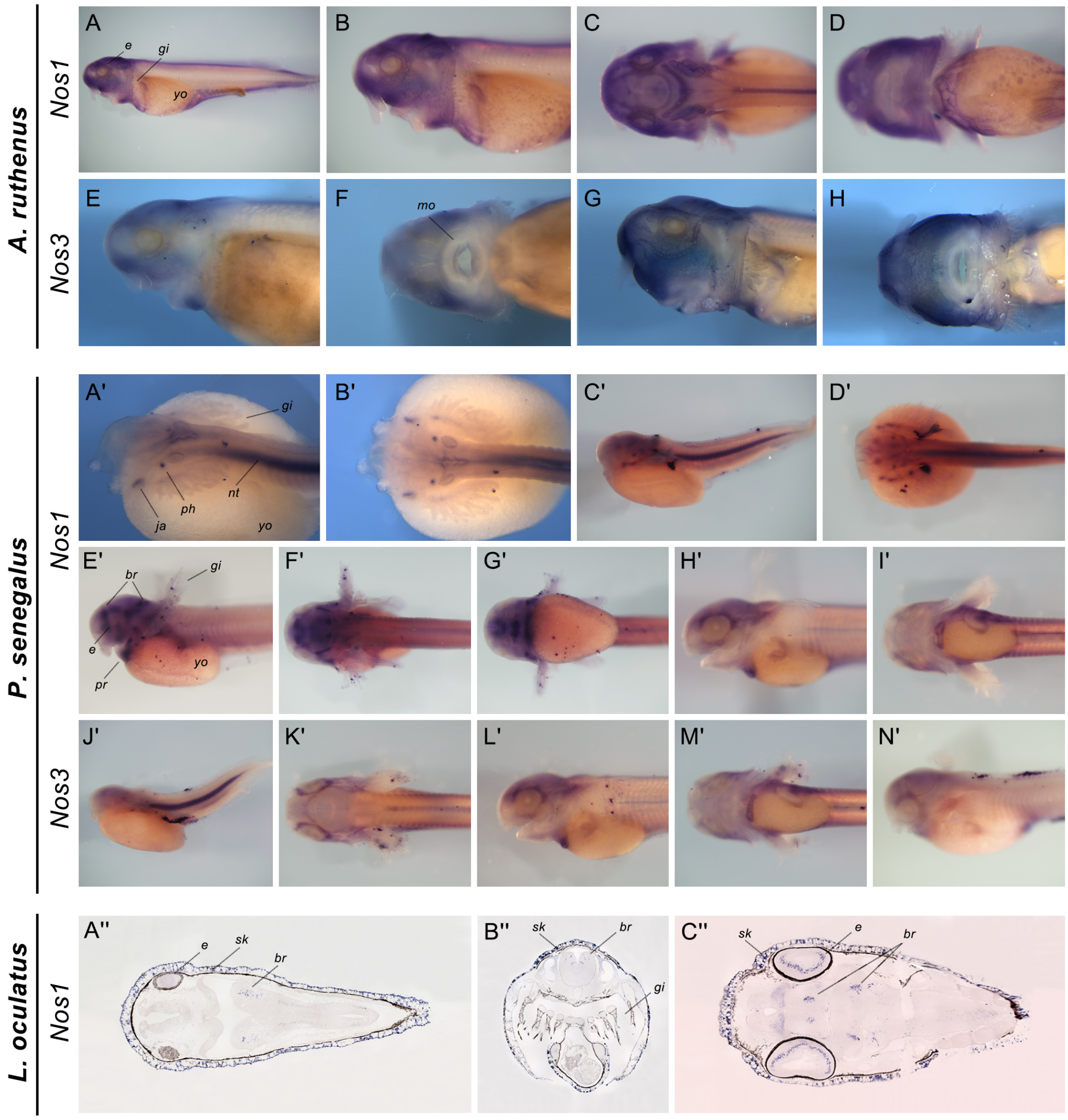


**Supplementary Figure 2.** Expression patterns of remaining *nos* genes in non-teleost fishes. *Acipenser ruthenus nos1* is expressed mainly in the head region, with intense staining in the brain (B-C), eye (C), and skin (A-D); *nos3* is diffusely expressed in the head skin (E-H). *Polypterus senegalus*, both *nos1* and *nos3* are expressed in the notochord at early developmental stages (A'-D', J'), and in the pericardial region (G'-I', L'-M'). Early in the development *nos1* is also expressed in the jaw joint area and pharynx (A'-B'), while later in the brain (E'-G'); *nos3* is expressed in the eye (H'). *Lepisosteus oculatus* (histological sections): *nos1* is expressed in several areas of the developing brain (A''-C''), eye (C''), and skin (A'', B'', C''). No expression was obtained for *nos2* during development. Abbreviations: br, brain; e, eye; gi, gill; ja, jaw joint area; mo, mouth; nt, notochord; ph, pharynx area; pr, pericardial region; sk, skin; yo, yolk.

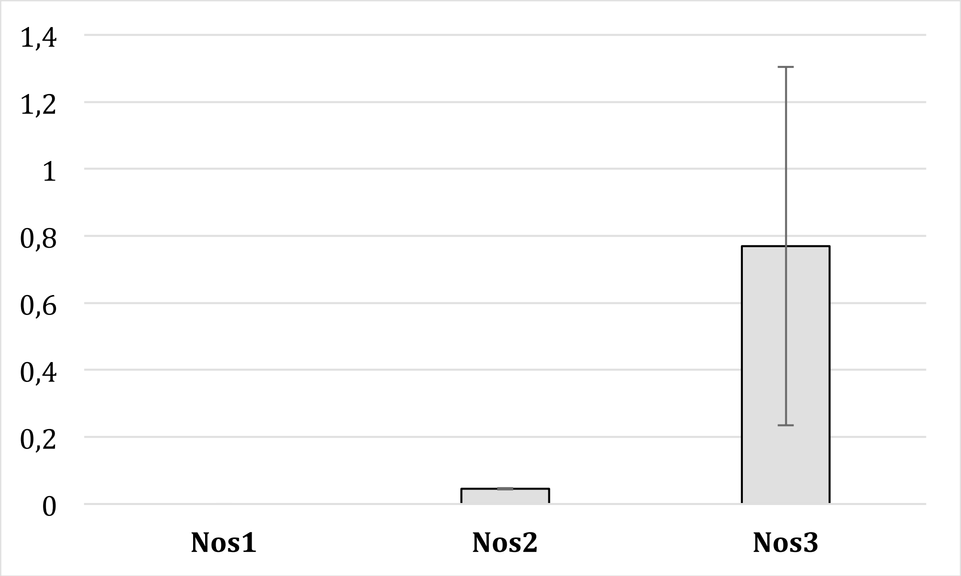


**Supplementary Figure 3.** Quantitative real-time PCR analysis of *nos* genes expression of axolotl *Ambystoma mexicanum* gills. Target gene expression was normalized to *atpf51* as a reference gene. Relative expression values were calculated using the ΔΔCT method. Error bars show the standard deviation of biological replicates. Expression estimates for the two *nos3* biological replicates were approximately 5x and 28x higher than the highest *nos2* estimate.

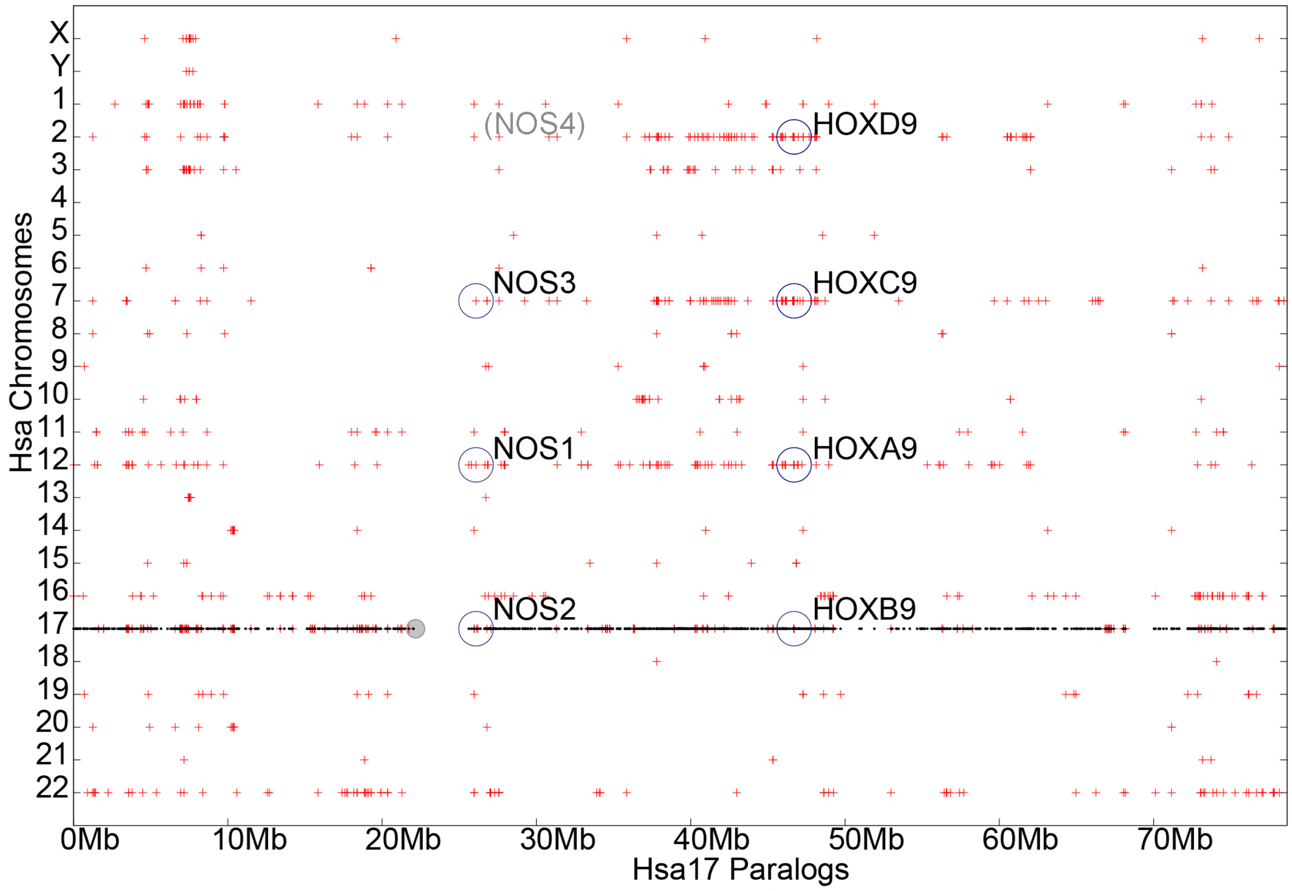


**Supplementary Figure 4.** Dotplot analysis showing syntenic conservation between human *NOS* genes and *HOX* clusters. Paralogs of human genes are plotted throughout the 23 chromosomes on Y-axis, within a 10 Mb-window, and are represented with red crosses. Dotplot shows syntenic conservation between four chromosomes harbouring *NOS* and *HOX* cluster genes. *NOS1* and *HOXA* cluster are on chromosome 12, *NOS2* and *HOXB* cluster are on chromosome 17, *NOS3* and *HOXC* cluster are on chromosome 7. According to the 2R hypothesis, a fourth *NOS* gene lost during evolution (*NOS4*) should have been positioned on chromosome 2 as the *HOXD* cluster. ENSG00000170689 (HOXB9) on Hsa17; ENSG00000128709 (HOXD9) on Hsa2; ENSG00000180806 (HOXC9) on Hsa12; ENSG00000078399 (HOXA9) on Hsa7; ENSG00000007171 (NOS2) on Hsa17; ENSG00000089250 (NOS1) on Hsa12; ENSG00000164867 (NOS3) on Hsa7.

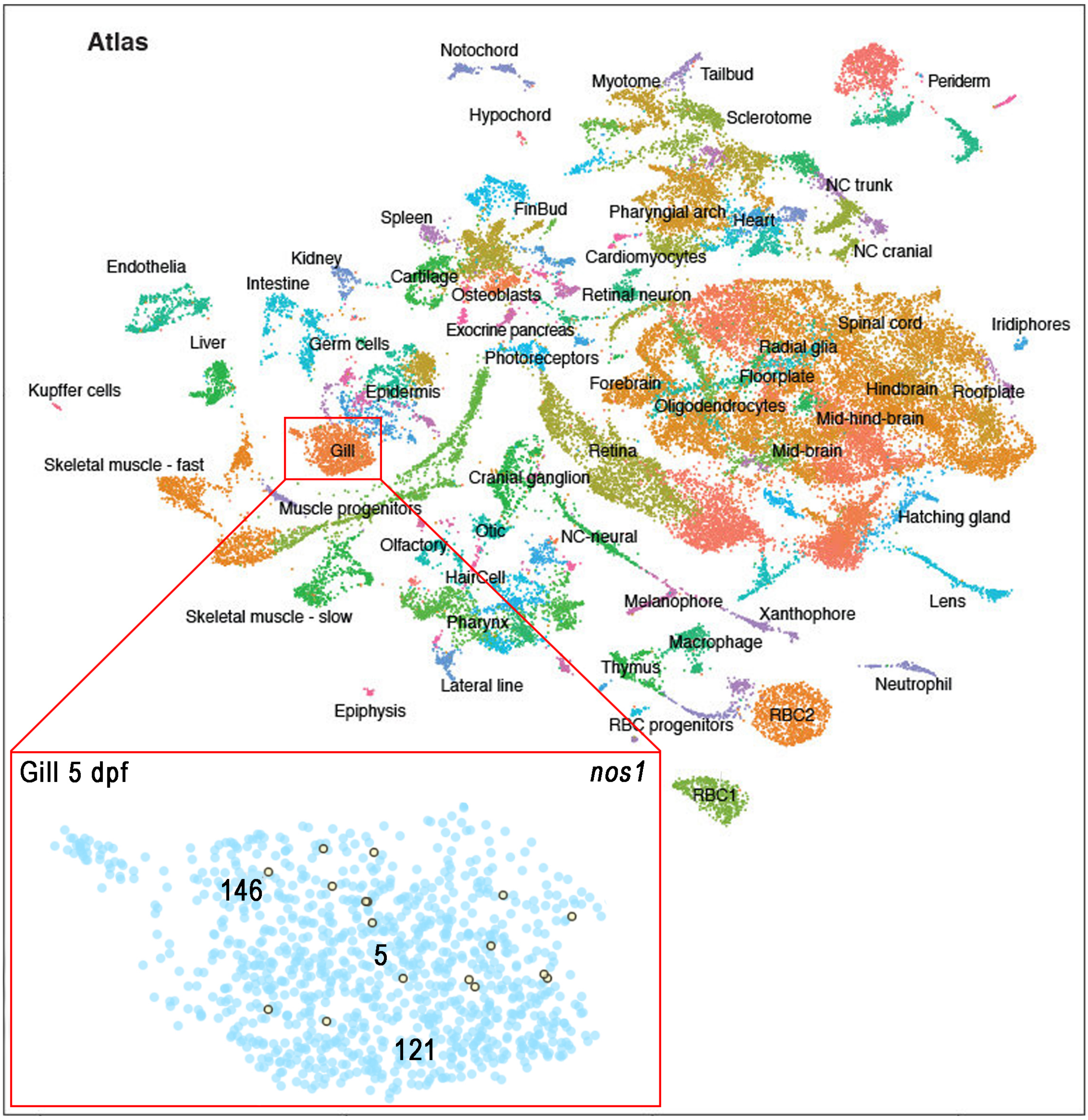


**Supplementary Figure 5.** Cell types clustering from single cell RNA-seq (scRNA-seq) atlas of zebrafish embryos during organogenesis (Miller lab, University of Oregon, Institute of Neuroscience, Eugene, Oregon). Colours correspond to annotated cell clusters. In the square are highlighted the cell clusters 5, 121 and 146 representing gill at 5 dpf and the specific *nos1*-expressing cells are shown in yellow. Data are obtained from the UCSC Cell Browser (http://zebrafish-dev.cells.ucsc.edu). Image modified from Farnsworth *et al.* (2020) [49].

**Supplementary File 1**

Alignment in FASTA format of the Nos protein sequences used to generate the phylogenetic tree of Figure 1.

**Supplementary Table 1**

Accession numbers of Nos proteins used in the phylogenetic tree of Figure 1.

| **Group/Species** | **Protein** | **Accession number** | **Database/Publication** |
|---|---|---|---|
| **Cephalochordates** | | | |
| *Branchiostoma lanceolatum* | NosA | MW652297 | NCBI GenBank |
| | NosB | MW652298 | NCBI GenBank |
| | NosC | MW652299, MW652300 | NCBI GenBank |
| **Tunicates** | | | |
| *Ciona robusta* | NosA | XP_002120267 | NCBI GenBank |
| **Cyclostomes** | | | |
| *Eptatretus burgeri* | Nos | MW652301 | This study |
| *Lethenteron camtschaticum* | NosA | MW652302 | This study |
| | NosB | MW652303 | This study |
| *Petromyzon marinus* | NosA | MW652304 | This study |
| | NosB | MW652305 | This study |
| **Chondrichthyes** | | | |
| *Callorhinchus milii* | Nos1 | XP_007900533 | NCBI GenBank |
| | Nos2 | XP_007898890 | NCBI GenBank |
| *Chiloscyllium punctatum* | Nos1 | GCC38273 | NCBI GenBank |
| | Nos2 | GCC31240 | NCBI GenBank |
| *Rhincodon typus* | Nos1 | XP_020369339 (corrected with Wise2) | NCBI GenBank |

| | | | |
|---|---|---|---|
| | Nos2 | XP_020382727 | NCBI GenBank |
| *Scyliorhinus torazame* | Nos1 | MW652306 | This study |
| | Nos2 | MW652307 | This study |
| | Nos3 | MW652308 | This study |
| **Acipenseriformes** | | | |
| *Acipenser ruthenus* | Nos2 | RXM97290 | NCBI GenBank |
| | Nos3 | RXM30658 | NCBI GenBank |
| **Polypteriformes** | | | |
| *Erpetoichthys calabaricus* | Nos1 | XP_028680314 | NCBI GenBank |
| | Nos2 | XP_028662939 | NCBI GenBank |
| | Nos3 | XP_028673556 | NCBI GenBank |
| *Polypterus senegalus* | Nos1 | MW574638 | NCBI GenBank |
| | Nos2 | MW574639 | NCBI GenBank |
| **Holosteans** | | | |
| *Amia calva* | Nos1 | | Thompson et al., 2020 |
| | Nos2 | | Thompson et al., 2020 |
| | Nos3 | | Thompson et al., 2020 |
| *Lepisosteus oculatus* | Nos1 | XP_015221897 | NCBI GenBank |
| | Nos2 | XP_006640988.2 | NCBI GenBank |
| | Nos3 | XP_015212637 | NCBI GenBank |
| **Teleosts** | | | |
| *Astyanax mexicanus* | Nos1 | XP_022523017 | NCBI GenBank |
| | Nos2.1 | XP_022536402 | NCBI GenBank |
| | Nos2.2 | XP_022536246 | NCBI GenBank |
| *Carassius auratus* | Nos1 | ENSCARP00000089103 | Ensembl |
| | Nos2a | ENSCARP00000054493 | Ensembl |
| | Nos2ba | ENSCARP00000115359 | Ensembl |
| | Nos2bb | ENSCARP00000111052 | Ensembl |

| | | | |
|---|---|---|---|
| *Clupea harengus* | Nos1 | XP_031426620 | NCBI GenBank |
| | Nos2.1 | XP_031428742 | NCBI GenBank |
| | Nos2.2 | XP_031428744 | NCBI GenBank |
| *Cynoglossus semilaevis* | Nos1 | XP_024909047 | NCBI GenBank |
| *Cyprinus carpio* | Nos1 | XP_018949869 | NCBI GenBank |
| | Nos2a | ENSCCRP00015121270 | Ensembl |
| | Nos2ba | ENSCCRP00015075878 | Ensembl |
| | Nos2bb | ENSCCRP00015045954 | Ensembl |
| *Danio rerio* | Nos1 | XP_005165110 | NCBI GenBank |
| | Nos2a | XP_005165353 | NCBI GenBank |
| | Nos2b | ABY51620 | NCBI GenBank |
| *Electrophorus electricus* | Nos1 | XP_026866423 | NCBI GenBank |
| | Nos2.1 | ENSEEEP00000042882 | Ensembl |
| | Nos2.2 | ENSEEEP00000017604 | Ensembl |
| *Esox lucius* | Nos1 | XP_010903911 | NCBI GenBank |
| | Nos2.1 | XP_012989745 | NCBI GenBank |
| | Nos2.2 | ENSELUP00000016280 | Ensembl |
| *Fundulus heteroclitus* | Nos1 | AAS21300.2 | NCBI GenBank |
| *Gadus morhua* | Nos1 | ENSGMOG00000014839 (annotation corrected with Wise2; partial sequence) | Ensembl |
| *Gasterosteus aculeatus* | Nos1 | ENSGACP00000018746 (annotation corrected with Wise2) | Ensembl |
| *Hippocampus comes* | Nos1 | XP_019741203 | NCBI GenBank |
| *Ictalurus punctatus* | Nos1 | XP_017306753 | NCBI GenBank |
| | Nos2.1 | XP_017347206 | NCBI GenBank |
| | Nos2.2 | ENSIPUP00000022197 | Ensembl |
| *Mastacembelus armatus* | Nos1 | XP_026177704 | NCBI GenBank |
| *Maylandia zebra* | Nos1 | XP_004566752 | NCBI GenBank |
| *Megalops cyprinoides* | Nos1 | XP_036382379 | NCBI GenBank |
| | Nos2 | XP_036392410 | NCBI GenBank |

| | | | |
|---|---|---|---|
| *Monopterus albus* | Nos1 | XP_020453884 | NCBI GenBank |
| *Neolamprologus brichardi* | Nos1 | XP_006803925 | NCBI GenBank |
| *Nothobranchius furzeri* | Nos1 | XP_015816026 | NCBI GenBank |
| *Oncorhynchus mykiss* | Nos1 | CDQ73326 | NCBI GenBank |
| | Nos2α | NP_001117831 | NCBI GenBank |
| | Nos2β | XP_021442676 | NCBI GenBank |
| *Oreochromis niloticus* | Nos1 | XP_013132314 | NCBI GenBank |
| *Oryzias latipes* | Nos1 | NP_001098325 | NCBI GenBank |
| *Paramormyrops kingsleyae* | Nos1 | XP_023684741 | NCBI GenBank |
| | Nos2 | XP_023692108 | NCBI GenBank |
| | Nos3 | XP_023681440 | NCBI GenBank |
| *Poecilia reticulata* | Nos1 | XM_017306782 | NCBI GenBank |
| *Pundamilia nyererei* | Nos1 | XP_005743120 | NCBI GenBank |
| *Salmo salar* | Nos1 | XP_014015466 | NCBI GenBank |
| | Nos2α | XP_014070450 | NCBI GenBank |
| | Nos2β | ENSSSAP00000062959 | Ensembl |
| *Scleropages formosus* | Nos1 | XP_018596057 | NCBI GenBank |
| | Nos2 | XP_018604029 | NCBI GenBank |
| *Scophthalmus maximus* | Nos1 | AWP07566 | NCBI GenBank |
| *Sciaenops ocellatus* | Nos1 | ACU98970 | NCBI GenBank |
| *Sinocyclocheilus anshuiensis* | Nos1 | XP_016362919 | NCBI GenBank |
| | Nos2a | ENSSANP00000063083 | Ensembl |
| | Nos2ba | ENSSANP00000007908 | Ensembl |
| | Nos2bb | ENSSANP00000004240 | Ensembl |
| *Stegastes partitus* | Nos1 | XP_008290591 | NCBI GenBank |
| *Takifugu rubripes* | Nos1 | AAL82736 | NCBI GenBank |
| *Tetraodon nigroviridis* | Nos1 | CAG08158 | NCBI GenBank |
| *Xiphophorus maculatus* | Nos1 | XP_005803890 | NCBI GenBank |
| **Coelacanthiformes** | | | |
| *Latimeria chalumnae* | Nos1 | XP_006008471 | NCBI GenBank |

| | | | |
|---|---|---|---|
| | Nos2 | XP_014346908 | NCBI GenBank |
| **Tetrapods** | | | |
| *Ambystoma mexicanum* | Nos1 | AMEXTC_0340000014481 | axolotl-omics.org |
| | Nos2 | AMEXTC_0340000001285+ AMEXTC_0340000212001 | axolotl-omics.org |
| | Nos3 | AMEXTC_0340000028637 | axolotl-omics.org |
| *Anolis carolinensis* | Nos1 | XP_008111929 | NCBI GenBank |
| | Nos2 | XP_003228867.2 | NCBI GenBank |
| | Nos3 | XP_008121260 | NCBI GenBank |
| *Homo sapiens* | NOS1 | AAB49040 | NCBI GenBank |
| | NOS2 | AAB60366 | NCBI GenBank |
| | NOS3 | AAM74944 | NCBI GenBank |
| *Xenopus tropicalis* | Nos1 | XP_004910558 | NCBI GenBank |
| | Nos2 | XP_002935342.2 | NCBI GenBank |
| | Nos3 | NP_001243158 | NCBI GenBank |

**Supplementary Table 2**

List of primers used for PCR amplification to prepare *nos* RNA probes and primers used for the qPCR analysis in axolotl *Ambystoma mexicanum.*

| ***In situ* hybridization primers** | | | |
|---|---|---|---|
| **Species** | **Gene** | **Sequence (5' > 3')** | **Riboprobe size** |
| *Lathenteron camtschaticum* | *nosA (3'UTR)* | *Fw* - GTAAACACGAATCTTCGCCT | 962 bp |
| | | *Rv* - ATGACAAATAACACACACCTAC | |
| | *nosB (3'UTR)* | *Fw* - GGGCAATATCATGGGCAAGC | 903 bp |
| | | *Rv* - GGATAACCACTCTTGACTTC | |
| *Lepisosteus oculatus* | *nos1* | *Fw* - CGACATGAGCTATGAGAGCG | 943 bp |
| | | *Rv* - GTCCACATGTGCCTTAGAGC | |
| | *nos2 (3'UTR)* | *Fw* - ATTCGGAGCAGTGTTTCAGA | 907 bp |
| | | *Rv* - CCCTTAGACCGTCGGTATTC | |
| | *nos3 (3'UTR)* | *Fw* - CACCTCCCAGATACGCTACC | 661 bp |
| | | *Rv* - CCTTCACTCCATATTCAGCC | |
| *Acipenser ruthenus* | *nos1* | *Fw* - GAAYTKTACACMGCMTACTC | 365 bp |
| | | *Rv* - AGAAAACCTCATCWSAGTCCT | |
| | *nos2* | *Fw* - TTSAGGTACTGTGTRTTTGG | 653 bp |
| | | *Rw* - AGGAAGTAGGTSAGGGYCTG | |
| | *nos3* | *Fw* - TGGGTAYWGCMWRCCAGATG | 645 bp |
| | | *Rv* - TAGTTVAGCAWYTCCTGGTG | |
| *Polypterus senegalus* | *nos1* | *Fw* - ATTTCCWCARCGRACAGATG | 440 bp |
| | | *Rv* - CAAKRATGTTATARCGAGAG | |
| | *nos2* | *Fw* - TTSAGGTACTGTGTRTTTGG | 664 bp |
| | | *Rv* - YGGTGATGTCSAGGAAGTAG | |
| | *nos3* | *Fw* - RTYGGMTYGAGGAACCTCTG | 298 bp |
| | | *Rv* - GGAAGACAGGVGTAABACTG | |
| *Scyliorhinus torazame* | *nos1 (3'UTR)* | *Fw* - GCATGCATCTCTTCCACCCT | 1041 bp |
| | | *Rv* - TGCCGCCACGTATCGATAAA | |
| | *nos2* | *Fw* - GGTGGAGCTCGGTCAATTCA | |
| | | *Rv* - GGTGGGGTTGTGGAAGGATT | 574 bp |

| | | | |
|---|---|---|---|
| | *nos3* | *Fw* - CGATGAGGAATCTTTAGGAA | 568 bp |
| | | *Rv* - CATATGTGTCCCCTGACTCA | |
| **qPCR primers** | | | |
| **Species** | **Gene** | **Sequence (5'>3')** | **Amplicon size** |
| *Ambystoma mexicanum* | *nos1* | *Fw* - GTGTCCAGTCTACTCCTGGAGATT | 140 bp |
| | | *Rv* - TTTCTTTGCGACTTCCTCTAAGAT | |
| | *nos2* | *Fw* - GAAGTATCACCCCCGTATTTCACC | 247 bp |
| | | *Rv* - GTTTCAGATTTCCCAGTTTCAGTT | |
| | *nos3* | *Fw* - GGAAATATAAGAGCAGCCATCACT | 100 bp |
| | | *Rv* - GGTAGCCAGCATAACGAAAGTATT | |
